# APOE4 Genotype is Associated with Reduced Cortical VEGFR2 (*KDR*) Transcript Levels Independent of Endothelial Abundance: An AMP-AD RNA-seq Pilot Study

**DOI:** 10.64898/2026.03.01.708815

**Authors:** K Laing, A Montagne

## Abstract

**Background:** Apolipoprotein E4 (*APOE4*) is the strongest common genetic risk factor for late-onset Alzheimer’s disease and has been implicated in cerebrovascular dysfunction. Vascular endothelial growth factor receptor 2 (VEGFR2), encoded by the gene *KDR*, is a key regulator of endothelial integrity and blood-brain barrier (BBB) maintenance; however, its relationship to APOE4 status in human brain tissue remains unclear.

**Methods:** Bulk RNA-sequencing data from 162 donors (n = 633 cortical samples) from the AMP-AD MSSB cohort were analysed using linear mixed-effects models with donor-level random intercepts. Models were adjusted for tissue region, age at death, sex, RNA integrity (RIN), and post-mortem interval. An endothelial composite score derived from key endothelial markers (*PECAM1, VWF, CLDN5, FLT1, TEK*) was included to account for endothelial signal abundance. Interaction models assessed modification by cortical region and neuropathological burden (Braak stage, CERAD score, plaque mean). A candidate panel of endothelial/BBB transcripts (*KDR, CAV1, MFSD2A, CLDN5, SLC2A1, OCLN, TJP1*) were also assessed in relation to APOE genotype.

**Results:** Bulk *KDR* expression varied significantly across cortical regions (p < 2×10^−16^) and was strongly correlated with endothelial signal abundance (r = 0.57, p < 2×10^−16^). *APOE4* status was not associated with endothelial composite score (p = 0.78). In models adjusting for tissue and technical covariates, but not endothelial signal, *APOE4* carriers showed a non-significant trend toward reduced *KDR* expression (β = −0.16, p = 0.064). After inclusion of the endothelial composite score, *APOE4* carrier status was significantly associated with reduced *KDR* expression (β = −0.17, p = 0.029), corresponding to an estimated 10-15% decrease in transcript levels. The direction of effect was consistent across cortical regions, with no evidence of *APOE4* × region interaction. No significant *APOE4* × neuropathology interactions were observed. Among candidate transcripts, only *KDR* and *TJP1* showed reduced expression, suggesting a selective rather than global transcriptional effect.

**Conclusions:** Bulk *KDR* transcript levels are strongly influenced by endothelial signal abundance; however, *APOE4* is associated with a modest but consistent reduction in VEGFR2 expression independent of endothelial signal abundance and neuropathological severity. These findings support a model in which *APOE4* contributes to vascular dysfunction through targeted endothelial alterations, rather than merely reflecting changes in endothelial cell proportion.

## Introduction

The ε4 allele of Apolipoprotein E (*APOE*) is a well-established genetic risk factor for late-onset Alzheimer’s disease (AD). The human APOE protein is a key regulator of lipid metabolism and exists in three common isoforms: E2, E3, and E4. Isoform structure is determined by two single-nucleotide polymorphisms (rs429358 and rs7412), which influence lipid-binding properties and receptor interactions with low density lipoproteins (LDL) and very low-density lipoproteins (VLDL) (1,2).

Growing preclinical and clinical evidence indicates that APOE4 carrier status contributes to cerebrovascular and BBB impairment (3,4). Many studies have focused on glial and mural components of the neurovascular unit, frequently describing a “leaky” BBB phenotype associated with increased matrix metalloproteinase activity and disrupted tight junction integrity (5–9). In a recent systematic review and meta-analysis of APOE target-replacement murine models, we identified convergent vascular phenotypes in APOE4, including reduced cerebral blood flow, altered vascular morphology, and increased BBB permeability (10). Three mechanistic pathways emerged: Cyclophilin A-NFκB-MMP9 activation, insulin resistance/PI3K/AKT-mTOR signalling dysregulation, and occludin phosphorylation linked to extracellular matrix remodelling.

Vascular endothelial growth factor receptor 2 (VEGFR2), encoded by the gene *KDR*, is a central regulator of endothelial function, angiogenic signalling, and BBB maintenance. Given its role in endothelial biology and its links to the pathways described above, we sought to determine whether VEGFR2 transcript levels are altered in *APOE4* carriers in human post-mortem brain tissue, and whether such effects are independent of endothelial signal abundance.

We hypothesised that *APOE4* carrier status is associated with reduced *VEGFR2* expression, reflecting endothelial functional alterations rather than changes in endothelial cell proportion.

## Methods

### AMP-AD Data Access

RNA-seq data were obtained from the AMP-AD RNA-seq Harmonization Study(11–13) via Synapse (accessed 25 February 2026). The full Mount Sinai Brain Bank (MSBB) dataset includes 315 donors; analyses were restricted to individuals with available *APOE* genotype and complete covariate data, resulting in 162 donors contributing 633 cortical samples. The MSSB cohort comprises bulk RNA-seq data from multiple cortical regions with corresponding neuropathological assessments, including Braak stage, CERAD score, and quantitative plaque burden. Regions analysed included parahippocampal gyrus (PHG), frontal pole (FP), inferior frontal gyrus (IFG), prefrontal cortex (PFC), and superior temporal gyrus (STG). Participants carrying an *APOE* ε2 allele (ε2/ε2, ε2/ε3, ε2/ε4) were excluded to isolate ε4-specific effects. Ages reported as “90+” in the original dataset were recoded as 90 years for statistical analyses.

### RNA-seq Processing

Gene-by-sample raw count matrices were downloaded from Synapse. Non-gene summary rows (e.g., alignment metrics) were removed prior to analysis. Ensembl gene identifiers were processed to remove version suffixes to ensure consistency across datasets. Counts were transformed to log2 counts per million (logCPM) using library-size normalisation with the ‘edgeR’ package in R.

### APOE Genotype

APOE genotyping was performed as described in the AMP-AD study details. DNA was extracted from peripheral blood mononuclear cells (PBMCs) or brain tissue. Genotyping targeted codon 112 (position 3937) and codon 158 (position 4075) of exon 4 of the *APOE* gene on chromosome 19 using high-throughput sequencing approaches (14). All *APOE* data were generated by Polymorphic DNA Technologies as part of a collaboration with Allan Roses and Zinfandel.

For this analysis, participants were classified as *APOE4* carriers (ε4 present: ε3/ε4 or ε4/ε4) versus ε3/ε3 non-carrier (reference group).

### Endothelial Signal Composite Score

An endothelial signal composite score was derived to estimate endothelial signal abundance within bulk RNA-seq samples. Endothelial markers included platelet endothelial cell adhesion molecule 1 (*PECAM1*; **ENSG00000261371**), von Willebrand factor (*VWF*; **ENSG00000110799**), claudin 5 (*CLDN5*; **ENSG00000184113**), fms-related tyrosine kinase 1 (*FLT1*(VEGFR1); **ENSG00000102755**), and TEK receptor tyrosine kinase (*TEK*; **ENSG00000120156**). For each gene, logCPM values were standardised (z-scored). The endothelial composite score was calculated as the mean of standardised expression values across these markers for each sample. This composite score was used as a proxy for endothelial signal in bulk tissue. Since VEGFR2 is primarily expressed in endothelial cells, models were adjusted for endothelial composite score to distinguish effects driven by cellular compositional (endothelial abundance) from per-cell transcriptional differences (15).

### Candidate Gene Analysis

Candidate endothelial/BBB transcripts (*KDR, CAV1, MFSD2A, CLDN5, SLC2A1, OCLN, TJP1*) were extracted from the logCPM-normalised RNA-seq matrix using Ensembl gene identifiers. For each gene, linear mixed-effects models were fitted to assess the association with *APOE4* carrier status, adjusting for endothelial composite score, tissue (region), age at death, sex, RIN, post-mortem interval (PMI), sequencing batch, and a donor-level random intercept.

For exploratory visualisation, unsupervised dimensionality reduction (uniform manifold approximation and projection; UMAP) was applied to scaled gene expression values. To assess genotype-associated structure independent of dominant confounders, gene expression values were additionally adjusted for tissue and technical covariates prior to embedding.

### Statistical Analysis

All analyses were conducted in R (RStudio version 2024.12.1). Linear mixed-effects models were fitted using ‘lmer’ package. The primary model was specified as: KDR_logCPM ∼ *APOE4* + tissue + endothelial composite score + covariates + (1 | individual ID). Covariates included APOE4 carrier status (ε4 present vs ε3/ε3), tissue (region of interest), endothelial composite score, sex, RIN (provided by AMP-AD for the technical integrity of RNA processing), PMI, and sequence batch. Individual ID was included as a random intercept to account for non-independence of multiple regional samples per donor. Interaction models were used to assess potential modification by tissue, Braak stage, CERAD score, and plaque mean. Estimated marginal means were computed using the ‘emmeans’ package, and likelihood ratio tests were used to compare nested models where appropriate.

## Results

The MSBB cohort included 315 donors, of whom 162 met inclusion criteria following quality control and filtering procedures. The final analytic cohort was 61.1% female and predominantly White. Mean RNA Integrity Number (RIN) was 6.67, indicating acceptable RNA quality for transcriptomic analyses.

Notably, *APOE4* carriers demonstrated greater neuropathological burden compared to non-carriers, including higher Braak stage, CERAD scores, and plaque mean (**Table 1**). However, interaction analyses (see section below) indicated that genotype-associated reductions in VEGFR2 expression were independent of neuropathological stage.

**Table 1.** Cohort Demographics by *APOE4* Carrier Status.

| Characteristic | Non-Carriers (n = 104) | *APOE4* Carriers (n = 58) | Total (n = 162) | p-value |
| --- | --- | --- | --- | --- |
| **Sex, n (%)** |  |  |  | 0.23 |
| Female | 60 (57.7%) | 39 (67.2%) | 99 (61.1%) |  |
| **Ethnicity, n (%)** |  |  |  | 0.41 |
| White | 90 (86.5%) | 46 (79.3%) | 136 (84.0%) |  |
| Hispanic | 5 (4.8%) | 5 (8.6%) | 10 (6.2%) |  |
| Black | 8 (7.7%) | 7 (12.1%) | 15 (9.3%) |  |
| Asian | 1 (1.0%) | 0 (0%) | 1 (0.6%) |  |
| **Age at death, years (mean)** | 82.9 ± 7.5 | 82.9 ± 7.5 | 82.9 ± 7.5 | 0.959 |
| **Post-mortem interval (PMI)** | 429.84 | 417.64 | 425.48 | 0.52 |
| **RNA Integrity Number (RIN)** | 6.75 | 6.54 | 6.67 | 0.21 |
| **CERAD score (mean)** | 1.92 | 2.50 | 2.13 | 0.003 ** |
| **Braak stage (mean)** | 3.62 | 4.29 | 3.87 | 0.02 * |
| **Clinical Dementia Rating (CDR)** | 2.30 | 2.47 | 2.36 | 0.31 |
| **Plaque mean (quantitative)** | 7.23 | 10.31 | 8.35 | 0.01 * |
| ***APOE* genotype distribution** | E3/E3: 104 (100%) | E3/E4: 53 (91.4%)<br>E4/E4: 5 (8.6%) | — | — |

### Bulk *KDR* Expression Varies Across Cortical Regions

Bulk expression of *KDR* differed significantly across cortical regions (F(4,597) = 23.45, p < 2.2×10^−16^), demonstrating marked regional heterogeneity in the absence of endothelial adjustment (**Figure 1A**; **Table 2**).

**Figure 1.**
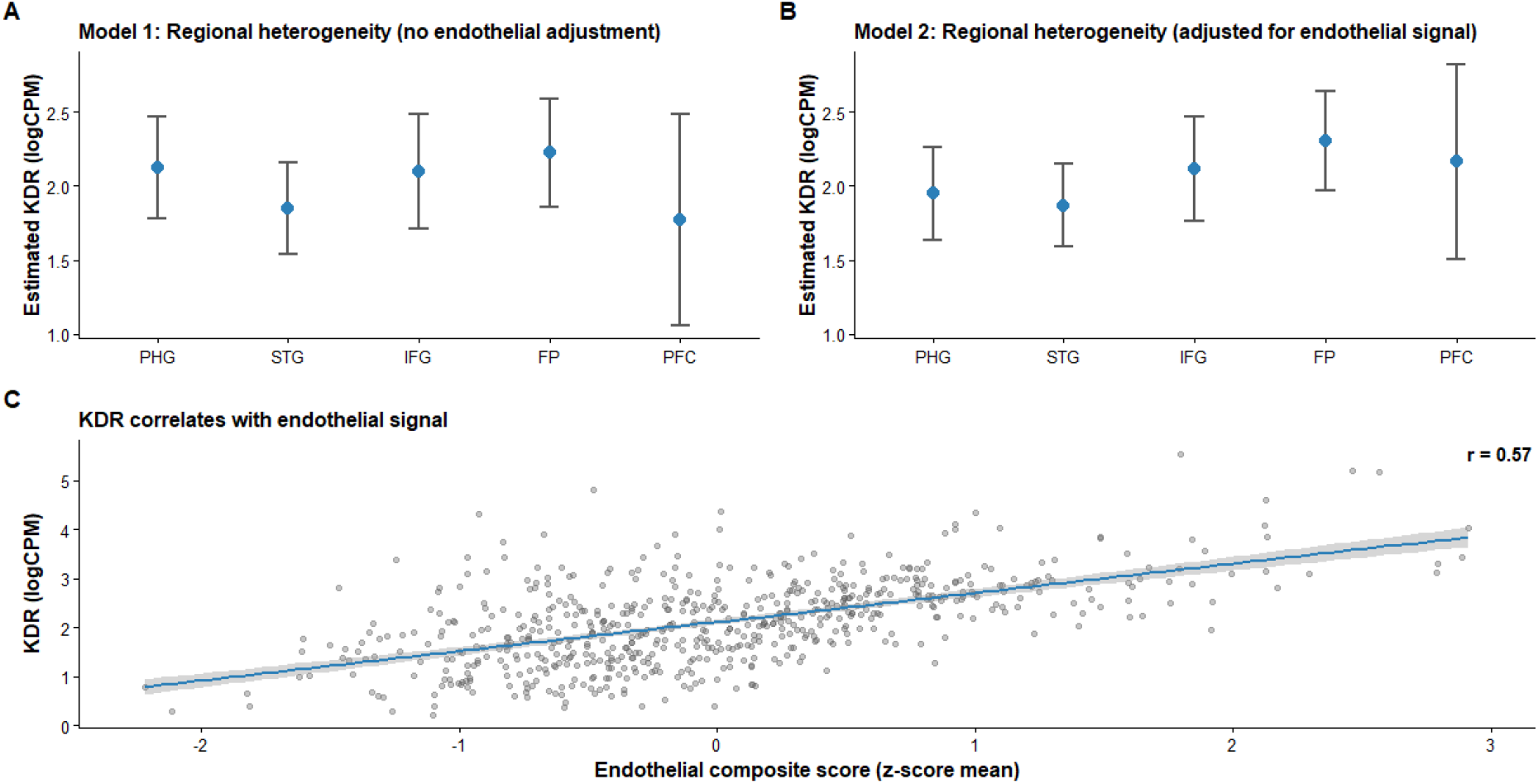
Regional heterogeneity and endothelial dependence of VEGFR2 (*KDR*) expression in human cortex. **(A)** Model-estimated marginal means (±95% CI) of bulk *KDR* expression (logCPM) across cortical regions without endothelial adjustment. **(B)** Model-estimated marginal means (±95% CI) of *KDR* expression after inclusion of an endothelial composite score derived from canonical endothelial markers (*PECAM1, VWF, CLDN5, FLT1*, and *TEK*). **(C)** Correlation between bulk *KDR* expression and endothelial composite score (Pearson r = 0.57, p < 0.001). *Abbreviations:* PHG, parahippocampal gyrus; STG, superior temporal gyrus; IFG, inferior frontal gyrus; FP, frontal pole; PFC, prefrontal cortex.

Inclusion of the endothelial composite substantially improved model fit and accounted for a large proportion of *KDR* variance (F(1,596) = 301.00, p < 2.2×10^−16^). Regional differences in VEGFR2 expression remained highly significant even after adjusting for endothelial signal (F(4,596) = 28.18, p < 2.2×10^−16^) (**Figure 1B**).

These findings indicate that while endothelial signal abundance strongly influences bulk *KDR* expression, it does not solely explain regional heterogeneity.

**Table 2.**
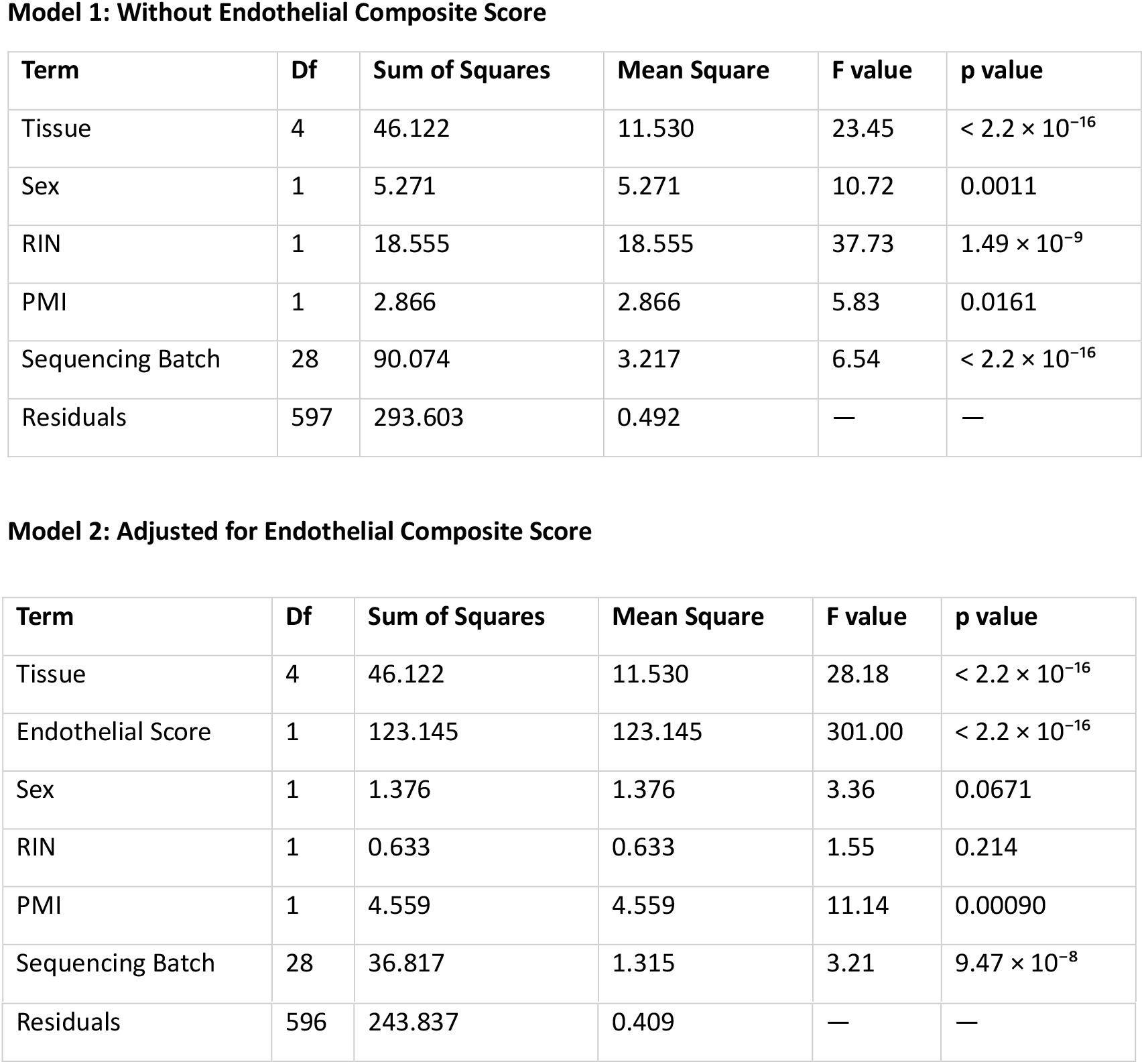
Analysis of Variance for VEGFR2 (*KDR*) Expression Across Cortical Regions. ANOVA results for VEGFR2 (*KDR*) bulk RNA-seq expression across cortical regions. Model 1 evaluates regional heterogeneity without endothelial adjustment. Model 2 includes an endothelial composite score derived from canonical endothelial markers (*PECAM1, VWF, CLDN5, FLT1*, and *TEK*). While endothelial signal strongly predicts VEGFR2 expression, regional differences remain significant after adjustment. *Abbreviations*: RIN = RNA Integrity Number; PMI = post-mortem interval.

### VEGFR2 Expression Strongly Correlates with Endothelial Signal

A composite endothelial signal score was derived from canonical endothelial markers (*PECAM1, VWF, CLDN5, FLT1, TEK*) (**Figure 1C**).

Bulk *KDR* expression demonstrated a strong positive correlation with the endothelial composite score (Pearson r = 0.57, p < 0.001), indicating that approximately 32% of the variance in *KDR* expression is explained by endothelial signal intensity.

These findings confirm that variation in bulk *KDR* expression largely reflects endothelial signal abundance within cortical tissue.

### APOE4 Does Not Alter Endothelial Signal

Endothelial composite scores did not differ between *APOE4* carriers and non-carriers (β ≈ 0.02, p = 0.78; **Figure 2**).

**Figure 2.**
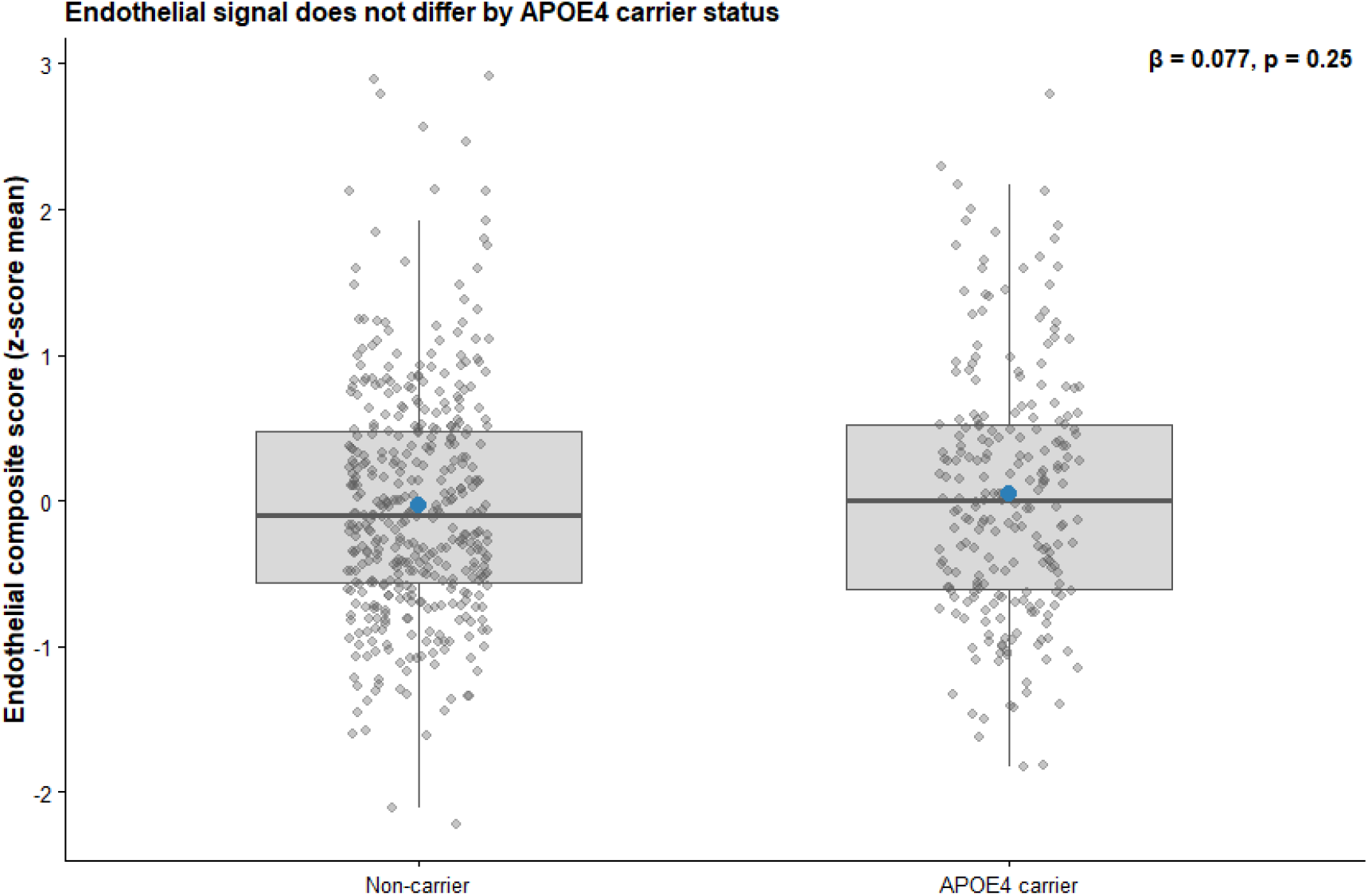
*APOE4* carrier status does not alter endothelial signal in bulk cortical RNA-seq. Endothelial composite scores were derived from canonical endothelial markers (*PECAM1, VWF, CLDN5, FLT1, TEK*) and calculated as the mean z-scored expression per sample. Points represent individual samples; boxplots show median and interquartile range; blue points denote group means. These findings indicate that *APOE4* genotype is not associated with reduced endothelial signal abundance in bulk cortical tissue.

Thus, *APOE4* carrier status is not associated with altered endothelial signal in bulk cortical tissue in this cohort.

### *APOE4* Is Associated with a Modest Global Reduction in *KDR* After Adjustment for Endothelial Signal

When examining bulk *KDR* expression between *APOE4* carriers and non-carriers, mixed-effects models adjusting for tissue and technical covariates—but not endothelial composite score— demonstrated a non-significant trend toward reduced *KDR* expression in *APOE4* carriers (β = −0.16, p = 0.064; **Figure 3A**).

**Figure 3.**
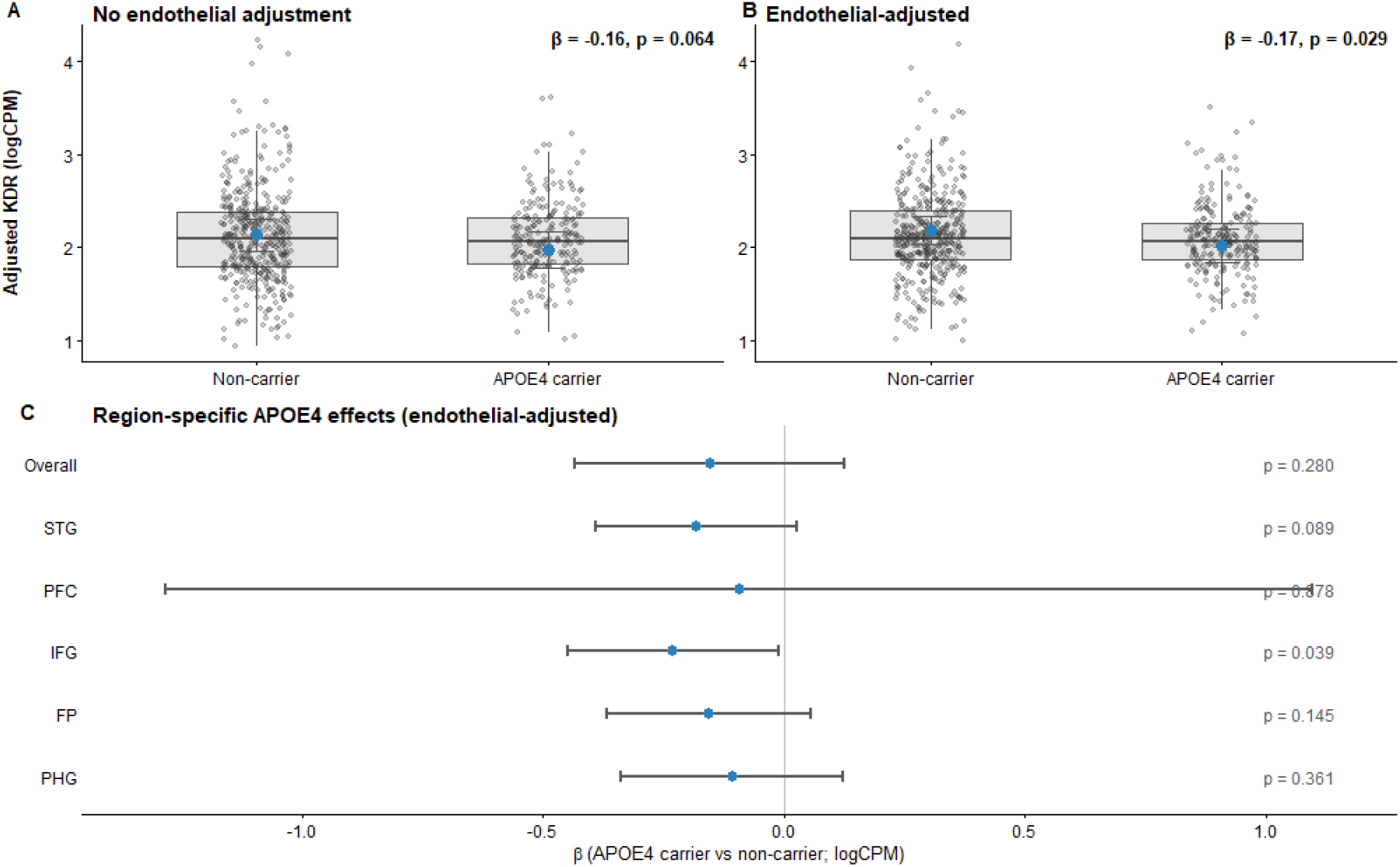
*APOE4* is associated with reduced VEGFR2 (*KDR*) expression after adjustment for endothelial signal. **(A)** Linear mixed-effects model of *KDR* logCPM comparing *APOE4* carriers vs non-carriers, adjusting for tissue, age at death, sex, RIN, PMI, sequencing batch, and a donor-level random intercept, without the endothelial composite score. **(B)** The same model including the endothelial composite score (*PECAM1, VWF, CLDN5, FLT1, TEK*), revealing a significant *APOE4*-associated reduction in *KDR* expression after accounting for endothelial signal. Points show covariate-adjusted individual values; blue points and whiskers denote model-estimated marginal means ± 95% confidence intervals. **(C)** Forest plot of region-specific *APOE4* effects on *KDR* (β for *APOE4* carriers vs non-carriers; 95% CI) from the endothelial-adjusted mixed-effects model, showing a consistent direction of effect across cortical regions with no evidence of region-specific modification.

Following inclusion of the endothelial composite score, *APOE4* carrier status was significantly associated with reduced *KDR* expression (β ≈ −0.17, p ≈ 0.0295), corresponding to an 10-15% reduction in transcript abundance (**Figure 3B**).

The direction of effect was consistent across all cortical regions examined (**Figure 3C**). Although only the inferior frontal gyrus reached statistical significance (β = -0.23, p = 0.040), no *APOE4* x tissue interaction was observed, supporting a global rather than region-specific effect.

### APOE4 Effect is Not Modified by Neuropathological Burden

Interaction models assessing *APOE4* × Braak stage, *APOE4* × CERAD score, and *APOE4* × plaque mean revealed no significant genotype-pathology interactions.

These findings indicate that the APOE4-associated reduction in *KDR* transcript levels is independent of amyloid and tau pathology severity.

### *APOE4* Effects on Candidate Endothelial/BBB Transcripts

Assessment of additional endothelial/BBB transcripts demonstrated limited genotype-dependent differences beyond VEGFR2 (*KDR*). In linear mixed models adjusted for endothelial composition, *TJP1* (ZO-1) also showed reduced transcript levels in *APOE4* carriers (**Table 3**; **Figure 4A**). Other candidate markers (*CAV1, MFSD2A, CLDN5, SLC2A1, OCLN*) did not demonstrate significant APOE4-associated changes.

**Figure 4.**
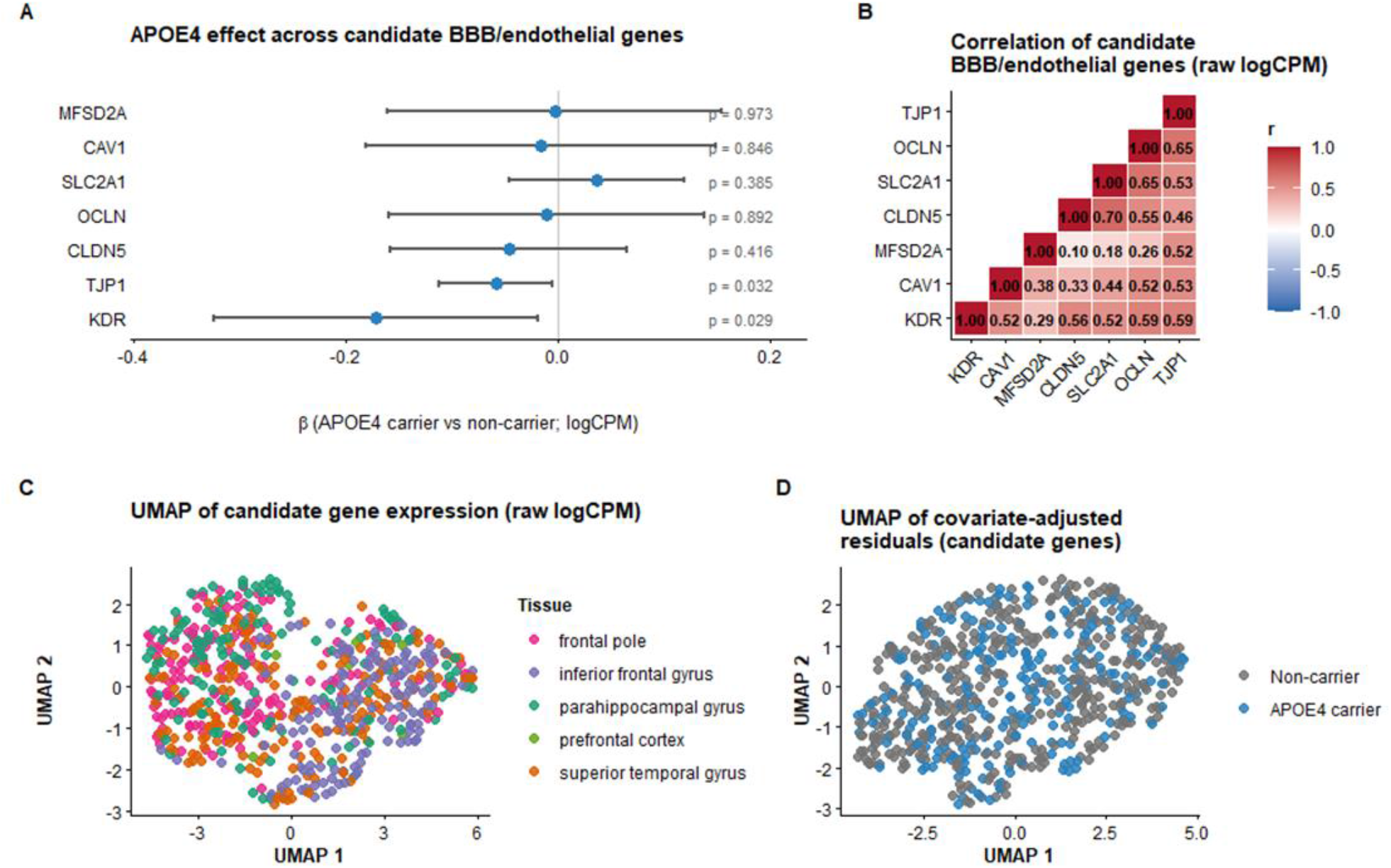
*APOE4* effects on candidate BBB/endothelial transcripts in bulk cortical RNA-seq. **(A)** Forest plot showing the estimated effect of *APOE4* carrier status on each candidate gene (β for *APOE4* carriers vs non-carriers; logCPM), derived from endothelial-adjusted linear mixed-effects models controlling for tissue, age at death, sex, RIN, PMI, sequencing batch, and donor-level random intercept. Only *KDR* and *TJP1* demonstrated nominally significant reductions in *APOE4* carriers; other canonical BBB/endothelial markers showed no genotype-associated differences. **(B)** Correlation matrix of raw logCPM expression values across candidate genes. Positive correlations were observed among endothelial/BBB-associated transcripts, consistent with shared vascular expression patterns in bulk cortical tissue. **(C)** UMAP embedding of raw logCPM values for the candidate gene panel across all samples. Samples cluster predominantly by cortical region (color-coded), indicating that regional differences represent the dominant source of variation in this gene set. **(D)** UMAP embedding of covariate-adjusted residual expression values (adjusted for tissue, age, sex, RIN, PMI, and sequencing batch). No clear separation by *APOE4* carrier status was observed, indicating that *APOE4* does not induce a broad transcriptional shift across endothelial/BBB markers.

Correlation analysis across candidate transcripts showed moderate-to-strong positive correlations (**Figure 4B**), consistent with shared vascular expression patterns in bulk tissue. Unsupervised embedding (UMAP) of raw expression values showed clear clustering by cortical region, indicating that regional identity is the dominant source of variation for these genes (**Figure 4C**).

After adjusting for tissue, technical covariates, and donor-level effects, no distinct clustering by *APOE4* carrier status was observed (**Figure 4D**). Together, these findings suggest that *APOE4* does not induce a broad endothelial transcriptional shift, but rather a selective reduction in specific targets, most notably *KDR* and, to a lesser extent, *TJP1*.

**Table 3.** *APOE4*-associated differences in endothelial and blood–brain barrier-related transcripts. Linear mixed-effects models estimating the association between *APOE4* carrier status and gene expression (logCPM), adjusting for endothelial composite score, tissue, age at death, sex, RIN, PMI, sequencing batch, and a donor-level random intercept.

| Gene | $\beta$ (*APOE4* carrier vs non-carrier) | SE | 95% CI | df | p-value |
| --- | --- | --- | --- | --- | --- |
| ***KDR*** | **−0.171** | 0.078 | −0.324 to −0.018 | 154.17 | **0.029** |
| *CAV1* | −0.016 | 0.084 | −0.181 to 0.148 | 156.65 | 0.846 |
| *MFSD2A* | −0.003 | 0.080 | −0.160 to 0.155 | 157.46 | 0.973 |
| *CLDN5* | −0.046 | 0.057 | −0.157 to 0.065 | 152.75 | 0.416 |
| *SLC2A1* | 0.037 | 0.042 | −0.046 to 0.119 | 162.10 | 0.385 |
| *OCLN* | −0.010 | 0.076 | −0.159 to 0.138 | 157.82 | 0.892 |
| ***TJP1*** | **−0.058** | 0.027 | −0.111 to −0.006 | 156.04 | **0.032** |

Overall, these findings demonstrate that bulk *KDR* expression is strongly driven by endothelial signal, while *APOE4* carrier status does not alter endothelial signal abundance. After accounting for endothelial signal, *APOE4* carriers exhibit a modest but consistent global reduction in *KDR* expression, which is not region-specific and is independent of neuropathological burden.

This pattern supports a model of *APOE4*-associated endothelial dysfunction characterised by targeted transcriptional changes rather than global endothelial loss.

## Discussion

This study assessed *KDR* transcript expression, encoding VEGFR2, in post-mortem cortical tissue from *APOE4* carriers and non-carriers. While altered APOE RNA and protein levels have been reported in AD (16), less is known about how *APOE* genotype influences downstream vascular-relevant transcriptional programs. Here, we demonstrate a modest but consistent reduction in *KDR* transcript levels in *APOE4* carriers that persists after adjusting for endothelial signal and technical covariates. Importantly, this finding suggests that *APOE4* is associated with endothelial functional alterations rather than changes in endothelial cell proportion.

Additionally, we show that the effect is not modified by neuropathological burden. Adjustment for CERAD score, plaque mean, and Braak stage did not attenuate the *APOE4*-*KDR* association, nor were genotype-by-pathology interactions observed. These findings indicate that *KDR* expression is unlikely to be secondary to amyloid or tau pathology, supporting a model in which *APOE4* contributes to vascular dysfunction through partially independent mechanisms.

Regional analysis demonstrated heterogeneity in bulk *KDR* transcript levels across cortical areas, consistent with previously reported regional differences in endothelial composition. Previous transcriptomic studies have shown that endothelial cells exhibit region-specific gene expression profiles in the human cortex(17). However, inclusion of an endothelial composite score in our models indicated that much of this regional variation is explained by endothelial signal. In contrast, the *APOE4*-associated reduction in *KDR* remained consistent across regions, suggesting a global genotype effect on endothelial transcriptional state.

Consistent with its well-established endothelial specificity (18,19), *KDR* expression correlated strongly with the endothelial composite score, irrespective of *APOE* genotype. Notably, *APOE4* carrier status was not associated with changes in the endothelial composite score, indicating that overall endothelial signal abundance is preserved in *APOE4* brains within this cohort. Taken together, these findings support a model in which *APOE4* selectively modulates endothelial gene expression—rather than inducing widespread endothelial loss—resulting in reduced *KDR* transcript levels within the endothelial compartment.

Further analyses examining *KDR* expression in relation to AD related neuropathology did not identify genotype-dependent modulation across stages of amyloid or tau burden. While prior studies have reported regional heterogeneity in endothelial responses to AD pathology(17), our results suggest that *APOE4*-associated *KDR* reduction occurs independently of these pathological processes, supporting the investigation of amyloid-independent vascular pathways.

Analysis of additional endothelial and BBB-related transcripts revealed limited genotype effects beyond *KDR*, with only a modest reduction observed in *TJP1* (ZO-1). Other canonical endothelial markers were not significantly altered. This argues against a broad transcriptional dysregulation of the endothelial/BBB program and instead supports a selective vulnerability of specific pathways, particularly VEGFR2 signalling.

While the current analyses demonstrate reduced *KDR* transcript levels, exploratory assessment of PI3K/AKT/mTOR pathway genes, mitochondrial markers, and oxidative stress-related transcripts did not reveal significant genotype-associated differences (**Supplementary Tables 1-3**). However, bulk RNA-seq is inherently limited in its ability to capture post-translational modifications and cell-type-specific signalling dynamics. It therefore remains possible that functional alterations in these pathways occur at the protein or phosphorylation level, which would not be detected in the present dataset. Future studies incorporating proteomics or phospho-proteomic approaches will be important to address these possibilities.

Several limitations should be considered. First, the use of bulk RNA-seq limits resolution of cell-type-specific effects and may obscure subtle endothelial or mural cell subtype differences. Second, the cross-sectional nature of post-mortem data precludes inference on temporal dynamics or causality. Third, although the endothelial composite score provides a useful proxy for endothelial signal, it cannot fully capture cellular heterogeneity within the vascular compartment.

Collectively, these findings highlight a robust and reproducible association between *APOE4* genotype and reduced *KDR* expression in the human cortex, independent of endothelial signal abundance and neuropathological burden. This positions VEGFR2 as a potential early node of vascular vulnerability in *APOE4* brains and supports a framework in which *APOE4* contributes to cerebrovascular dysfunction through targeted modulation of endothelial signalling pathways rather than global endothelial loss.

## Acknowledgements

The work of AM is supported by the UK Dementia Research Institute (award UKDRI-4209) through UK DRI Ltd, funded by the UK Medical Research Council (MRC), Alzheimer’s Society UK, Alzheimer’s Research UK, and the British Heart Foundation. AM also holds a UKRI MRC Career Development Award (MR/V032488/1) and a UK DRI Theme Funding Programme Award (DRI-TFP-2024-7). KKL received support from the University of Edinburgh Wellcome Trust Translational Neuroscience 4-year PhD Programme (Grant No. 110002 20132001 TBC 130956 00000000 10002374 000).

The results published here are in whole or in part based on data obtained from The AD Knowledge Portal (https://doi.org/10.7303/9618241). Data generation was supported by the following NIH grants: P30AG10161, P30AG72975, R01AG15819, R01AG17917, R01AG036836, U01AG46152, U01AG61356, U01AG046139, P50 AG016574, R01 AG032990, U01AG046139, R01AG018023, U01AG006576, U01AG006786, R01AG025711, R01AG017216, R01AG003949, R01NS080820, U24NS072026, P30AG19610, U01AG046170, RF1AG057440, and U24AG061340, and the Cure PSP, Mayo and Michael J Fox foundations, Arizona Department of Health Services and the Arizona Biomedical Research Commission. We thank the participants of the Religious Order Study and Memory and Aging projects for the generous donation, the Sun Health Research Institute Brain and Body Donation Program, the Mayo Clinic Brain Bank, and the Mount Sinai/JJ Peters VA Medical Center NIH Brain and Tissue Repository. Data and analysis contributing investigators include Nilüfer Ertekin-Taner, Steven Younkin (Mayo Clinic, Jacksonville, FL), Todd Golde (University of Florida), Nathan Price (Institute for Systems Biology), David Bennett, Christopher Gaiteri (Rush University), Philip De Jager (Columbia University), Bin Zhang, Eric Schadt, Michelle Ehrlich, Vahram Haroutunian, Sam Gandy (Icahn School of Medicine at Mount Sinai), Koichi Iijima (National Center for Geriatrics and Gerontology, Japan), Scott Noggle (New York Stem Cell Foundation), Lara Mangravite (Sage Bionetworks).

## References

1. Mahley RW. Apolipoprotein E: from cardiovascular disease to neurodegenerative disorders. J Mol Med. 2016 Jul 1;94(7):739–46. doi:10.1007/s00109-016-1427-y

2. Mahley RW. Apolipoprotein E: Cholesterol Transport Protein with Expanding Role in Cell Biology. Science. 1988 Apr 29;240(4852):622–30. doi:10.1126/science.3283935

3. Montagne A, Barnes SR, Sweeney MD, Halliday MR, Sagare AP, Zhao Z, et al. Blood-Brain Barrier Breakdown in the Aging Human Hippocampus. Neuron. 2015 Jan 21;85(2):296. doi:10.1016/j.neuron.2014.12.032 PubMed PMID: 25611508.

4. Montagne A, Nation DA, Sagare AP, Barisano G, Sweeney MD, Chakhoyan A, et al. APOE4 leads to blood-brain barrier dysfunction predicting cognitive decline. Nature. 2020 May;581(7806):71–6. doi:10.1038/s41586-020-2247-3 PubMed PMID: 32376954; PubMed Central PMCID: PMC7250000.

5. Jackson RJ, Meltzer JC, Nguyen H, Commins C, Bennett RE, Hudry E, et al. APOE4 derived from astrocytes leads to blood-brain barrier impairment. Brain. 2022;145(10):3582–93. doi:10.1093/brain/awab478

6. Bhattarai P, Yilmaz E, Cakir EÖ, Korkmaz HY, Lee AJ, Ma Y, et al. APOE-ε4-induced Fibronectin at the blood-brain barrier is a conserved pathological mediator of disrupted astrocyte-endothelia interaction in Alzheimer’s disease [Internet]. bioRxiv; 2025 [cited 2025 Jul 28]. p. 2025.01.24.634732. Available from: https://www.biorxiv.org/content/10.1101/2025.01.24.634732v1 doi:10.1101/2025.01.24.634732

7. Blanchard JW, Bula M, Davila-Velderrain J, Akay LA, Zhu L, Frank A, et al. Reconstruction of the Human Blood-Brain Barrier in vitro reveals a Pathogenic Mechanism of APOE4 in Pericytes. Nat Med. 2020 Jun;26(6):952–63. doi:10.1038/s41591-020-0886-4 PubMed PMID: 32514169; PubMed Central PMCID: PMC7704032.

8. Yamazaki Y, Shinohara M, Yamazaki A, Ren Y, Asmann YW, Kanekiyo T, et al. ApoE (Apolipoprotein E) in Brain Pericytes Regulates Endothelial Function in an Isoform-Dependent Manner by Modulating Basement Membrane Components. Arter Thromb Vasc Biol. 2020;40(1):128–44. doi:10.1161/ATVBAHA.119.313169

9. Yamazaki Y, Liu CC, Yamazaki A, Shue F, Martens YA, Chen Y, et al. Vascular ApoE4 Impairs Behavior by Modulating Gliovascular Function. Neuron. 2021;109(3):438-447.e6. doi:10.1016/j.neuron.2020.11.019

10. Laing K, Fialova N, Wardlaw J, Montagne A. Impact of Apolipoprotein E4 on Blood-Brain Barrier Integrity in Target Replacement Murine Models: A Systematic Review and Meta-Analysis [Internet]. Research Square; 2026 [cited 2026 Feb 27]. Available from: https://www.researchsquare.com/article/rs-8011437/v1 doi:10.21203/rs.3.rs-8011437/v1

11. Allen M, Carrasquillo MM, Funk C, Heavner BD, Zou F, Younkin CS, et al. Human whole genome genotype and transcriptome data for Alzheimer’s and other neurodegenerative diseases. Sci Data. 2016 Oct 11;3(1):160089. doi:10.1038/sdata.2016.89

12. Wang M, Beckmann ND, Roussos P, Wang E, Zhou X, Wang Q, et al. The Mount Sinai cohort of large-scale genomic, transcriptomic and proteomic data in Alzheimer’s disease. Sci Data. 2018 Sep 11;5(1):180185. doi:10.1038/sdata.2018.185

13. De Jager PL, Ma Y, McCabe C, Xu J, Vardarajan BN, Felsky D, et al. A multi-omic atlas of the human frontal cortex for aging and Alzheimer’s disease research. Sci Data. 2018 Aug 7;5(1):180142. doi:10.1038/sdata.2018.142

14. Yu L, Lutz MW, Wilson RS, Burns DK, Roses AD, Saunders AM, et al. TOMM40′523 variant and cognitive decline in older persons with APOE ε3/3 genotype. Neurology. 2017 Feb 14;88(7):661–8. doi:10.1212/WNL.0000000000003614 PubMed PMID: 28108637; PubMed Central PMCID: PMC5317377.

15. Farahbod M, Pavlidis P. Untangling the effects of cellular composition on coexpression analysis. Genome Res. 2020 Jun;30(6):849–59. doi:10.1101/gr.256735.119 PubMed PMID: 32580998; PubMed Central PMCID: PMC7370889.

16. Akram A, Schmeidler J, Katsel P, Hof PR, Haroutunian V. Association of ApoE and LRP mRNA levels with dementia and AD neuropathology. Neurobiol Aging. 2012 Mar;33(3):628.e1-628.e14. doi:10.1016/j.neurobiolaging.2011.04.010 PubMed PMID: 21676498; PubMed Central PMCID: PMC3234309.

17. Bryant A, Li Z, Jayakumar R, Serrano-Pozo A, Woost B, Hu M, et al. Endothelial Cells Are Heterogeneous in Different Brain Regions and Are Dramatically Altered in Alzheimer’s Disease. J Neurosci. 2023 Jun 14;43(24):4541–57. doi:10.1523/JNEUROSCI.0237-23.2023 PubMed PMID: 37208174; PubMed Central PMCID: PMC10278684.

18. Gampel A, Moss L, Jones MC, Brunton V, Norman JC, Mellor H. VEGF regulates the mobilization of VEGFR2/KDR from an intracellular endothelial storage compartment. Blood. 2006 Oct 15;108(8):2624–31. doi:10.1182/blood-2005-12-007484

19. Shah FH, Nam YS, Bang JY, Hwang IS, Kim DH, Ki M, et al. Targeting vascular endothelial growth receptor-2 (VEGFR-2): structural biology, functional insights, and therapeutic resistance. Arch Pharm Res. 2025;48(5):404–25. doi:10.1007/s12272-025-01545-1 PubMed PMID: 40341988; PubMed Central PMCID: PMC12106596.

