## Supplementary Tables for "APOE4 Genotype is Associated with Reduced Cortical VEGFR2 (*KDR*) Transcript Levels Independent of Endothelial Abundance: An AMP-AD RNA-seq Pilot Study"

**Supplementary Table 1. *APOE4* association with PI3K/AKT/mTOR pathway transcripts.** Linear mixed-effects models were fit for each gene: Gene ~ *APOE4* + endo_score + tissue + ageDeath + sex + RIN + pmi + sequencingBatch + (1|individualID).

| **Gene** | **β (*APOE4*)** | **SE** | **df** | **95% CI** | **p-value** | **FDR** |
| --- | --- | --- | --- | --- | --- | --- |
| *RPS6KB1* | 0.035 | 0.025 | 123 | −0.015 to 0.084 | 0.169 | 0.945 |
| *RPTOR* | 0.026 | 0.024 | 159 | −0.021 to 0.072 | 0.279 | 0.945 |
| *FOXO3* | 0.028 | 0.027 | 126 | −0.024 to 0.080 | 0.289 | 0.945 |
| *PIK3CA* | 0.027 | 0.029 | 131 | −0.029 to 0.084 | 0.349 | 0.945 |
| *PIK3CB* | 0.042 | 0.045 | 107 | −0.047 to 0.131 | 0.362 | 0.945 |
| *RICTOR* | 0.021 | 0.025 | 119 | −0.028 to 0.069 | 0.406 | 0.945 |
| *MTOR* | 0.019 | 0.028 | 110 | −0.037 to 0.074 | 0.510 | 0.945 |
| *PTEN* | 0.020 | 0.035 | 120 | −0.048 to 0.087 | 0.571 | 0.945 |
| *FOXO1* | −0.022 | 0.038 | 139 | −0.096 to 0.053 | 0.573 | 0.945 |
| *PDPK1* | 0.017 | 0.033 | 123 | −0.048 to 0.082 | 0.604 | 0.945 |
| *AKT1* | 0.010 | 0.020 | 135 | −0.029 to 0.048 | 0.626 | 0.945 |
| *AKT3* | 0.012 | 0.043 | 122 | −0.072 to 0.097 | 0.774 | 0.945 |
| *RHEB* | 0.013 | 0.051 | 123 | −0.087 to 0.113 | 0.799 | 0.945 |
| *TSC2* | −0.006 | 0.026 | 158 | −0.058 to 0.045 | 0.808 | 0.945 |
| *TSC1* | 0.005 | 0.019 | 131 | −0.032 to 0.041 | 0.811 | 0.945 |
| *PIK3CD* | −0.008 | 0.040 | 160 | −0.086 to 0.070 | 0.841 | 0.945 |
| *PIK3R2* | −0.008 | 0.041 | 162 | −0.088 to 0.072 | 0.850 | 0.945 |
| *AKT2* | 0.003 | 0.031 | 140 | −0.058 to 0.065 | 0.916 | 0.945 |
| *EIF4EBP1* | 0.006 | 0.063 | 159 | −0.117 to 0.129 | 0.924 | 0.945 |
| *PIK3R1* | 0.003 | 0.044 | 117 | −0.083 to 0.089 | 0.945 | 0.945 |

**Supplementary Table 2. *APOE4* association with mitochondrial transcriptional signature.** Linear mixed-effects model: mito_sig ~ *APOE4* + endo_score + tissue + ageDeath + sex + RIN + pmi + sequencingBatch + (1 | individualID). Random intercept: individualID.

Tissue and sequencing batch covariates included in model (coefficients not shown). Mitochondrial signature derived from mean z-scored expression of *NDUFS1*, *NDUFV1*, *SDHA*, *UQCRC1*, *COX4I1*, *ATP5F1A*, *TFAM*, and *PPARGC1A*.

| **Variable** | **β** | **SE** | **df** | **t** | **p-value** |
| --- | --- | --- | --- | --- | --- |
| ***APOE4*** | 0.037 | 0.088 | 104 | 0.42 | 0.678 |
| Endothelial composite | −0.069 | 0.045 | 594 | −1.54 | 0.125 |
| Age at death | −0.002 | 0.006 | 107 | −0.38 | 0.707 |
| Sex (male) | 0.105 | 0.093 | 105 | 1.12 | 0.265 |
| RIN | 0.334 | 0.025 | 584 | 13.23 | <0.001 |

**Supplementary Table 3. *APOE4* association with oxidative stress transcriptional signature.** Linear mixed-effects model: oxstress_sig ~ *APOE4* + endo_score + tissue + ageDeath + sex + RIN + pmi + sequencingBatch + (1 | individualID). Random intercept: individualID.

Tissue and sequencing batch covariates included in model (coefficients not shown). Oxidative stress signature derived from mean z-scored expression of *SOD1*, *SOD2*, *GPX1*, *CAT*, *NFE2L2*, *HMOX1*, *TXN*, and *PRDX1*.

| **Variable** | **β** | **SE** | **df** | **t** | **p-value** |
| --- | --- | --- | --- | --- | --- |
| ***APOE4*** | 0.047 | 0.069 | 136 | 0.69 | 0.493 |
| Endothelial composite | 0.288 | 0.036 | 594 | 8.07 | <0.001 |
| Age at death | 0.00008 | 0.00507 | 138 | 0.02 | 0.988 |
| Sex (male) | 0.069 | 0.073 | 137 | 0.95 | 0.346 |
| RIN | 0.233 | 0.020 | 589 | 11.59 | <0.001 |
